# Recombinant Protein IGF1-24 Stimulates Rat Cardiomyocytes Proliferation and Repairs Myocardial Injury

**DOI:** 10.1101/2020.03.04.976522

**Authors:** Xing Wang, Manxue Fu, Qian Yi, Jianguo Feng, Yi Liao, Xichao Xu, Ying Chen, Lu Zhang, Huifang Sun, Piaoyang Liu, Yuanyuan Liang, Liling Tang

**Affiliations:** Key Laboratory for Biorheological Science and Technology of Ministry of Education, College of Bioengineering, Chongqing University, Chongqing, China; Department of Physiology, School of Basic Medical Sciences, Southwest Medical University, Luzhou, Sichuan Province, China; Department of Anesthesiology, The Affiliated Hospital of Southwest Medical University, Luzhou, Sichuan Province, China; Department of Thoracic Surgery, Southwest Hospital, Army Medical University, Chongqing, China; College of Bioengineering, Chongqing University, Chongqing, China

**Keywords:** Recombinant Protein IGF1-24, Cardiomyocytes Proliferation, Myocardial Injury

## Abstract

Mammal cardiomyocytes lose their ability of regeneration shortly after birth. Reduced cardiomyocytes number caused by myocardial damage is unable to reverse in current clinical therapies. Therefore, it is important and urgent to find new approaches to stimulate cardiomyocytes regeneration. Here we design a recombinant protein IGF1-24 and show that it triggers cardiomyocytes proliferation in rat. 7 days after tail intravenous injection of IGF1-24, 6-7-weeks-old healthy rats showed marked improvements in cardiomyocytes proliferation. Next, we injured rats cardiac with isoproterenol and treated them with IGF1-24 injection. We found that it efficiently induced cell proliferation with significant improvements in heart histology. These results show that the recombinant IGF1-24 stimulates cardiomyocytes proliferation and can be used to achieve cardiac repair through stimulating endogenous cardiomyocyte proliferation in rats. The IGF 1-24 could be a prospective medicine to heart repair because it has high efficiency in triggering cell proliferation and it can be easily applied to heart by intravenous injection

## Introduction

Unlike organisms such as zebrafish[1], the regenerative ability of mammalian heart through cell proliferation is limited after birth[2–4]. Cardiomyocytes proliferation was almost undetectable in 7-days-old mice[5, 6] and rats[7]. Myocardial infarction, heart failure and other cardiac diseases reduce cardiomyocyte number. As a result, the cardiac function could not be recovered[8, 9]. Therefore, exploring new approaches and medicine to stimulate cardiomyocytes proliferation could be of great clinical significance.

Previous studies have shown that a few miRNAs can stimulate endogenous cardiomyocytes proliferation and cardiac regeneration in myocardial infarction mice[7, 10] and pigs[11]. Induction of cell-cycle genes including CDK1,CCNB, CDK4 and CCND94F induced cardiomyocyte proliferation and stimulated regeneration of the infarcted heart[12]. In addition, regulatory T cells (Tregs) promoted cardiomyocyte proliferation after myocardial infarction, and growth factors secreted by Tregs may play roles in the regeneration process[13]. Growth factors are key regulators of cell proliferation, differentiation and survival. And there are many successful clinical applications. Therefore, we attempt to explore a growth factor which can stimulate cardiomyctes proliferation and cardiac regeneration.

The mammalian insulin-like growth factor 1 (IGF1) plays important roles during mammal development in an autocrine/paracrine fashion. IGF 1 was able to increase mice growth[14]. IGF 1 deficiency exacerbated the pathogenesis of cerebral microhemorrhages, which is related to aging phenotype[15]. In addition, IGF1 treatment improved the hepataprotection in acute liver damage in mice[16]. However, there is no evidence showing that IGF1 has the ability to stimulate cardiomyocytes proliferation.

We noticed that the IGF 1Ec, a splicing variant of IGF 1,is produced when body is subjected to injury or mechanical load. Thus it is also named as mechanical growth factor (MGF)[17]. Related studies have reported that IGF1Ec can protect heart function [18]and nerve cells[19]. Still, no researches have shown that it can promote the proliferation of differentiated cardiomyocytes. Here we designed a recombinant protein IGF 1-24 based on IGF 1 and IGF 1 Ec. We treated the healthy adult rats with these proteins and found that the recombinant protein IGF1-24 effectively induced the proliferation of differentiated myocardial cells, the effect is 2.4 times higher than IGF 1Ec. IGF 1 has no effect on proliferation of cardiomyocytes (FigS1).

In this paper, we investigated the effects of IGF 1-24 on cardiomyocytes proliferation of rats. Results show that treatment of the IGF 1-24 efficiently induces cardiomyocyte proliferation in vivo. Tail intravenous injection of IGF1-24 resulted in improved cardiac regeneration after injury compared to controls.

## Results

### Design of IGF 1-24

IGF1-24 sequence was constructed by linking IGF1 and 24 amino acids which was cloned from C-terminal of IGF 1 Ec by three glycines as shown in our previous report[20](Fig1a). Then we expressed the recombinant protein in rosetta E.coil and purified it with Ni-Agaros (Fig1b). The recombinant protein IGF1-24 contains 97 amino acids with molecular weight of 10.67 Kd.

**Fig.1.**
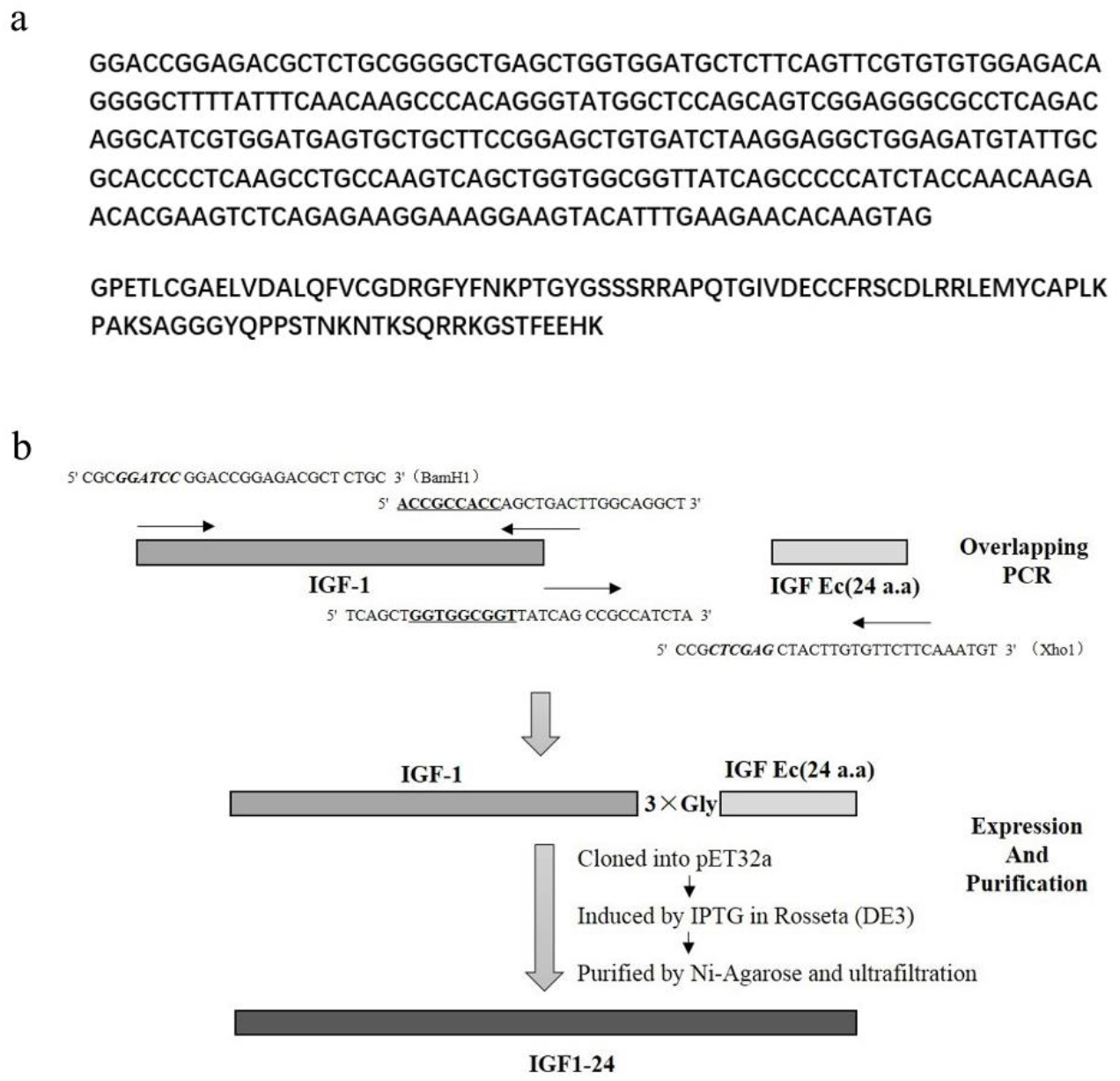
Structure of IGF1-24. **a,** Gene sequences and amino acid sequences of IGF1-24. **b,** Schematic diagram of IGF1-24 construction and expression.

### IGF1-24 stimulated full-differentiated cardiomyocytes proliferation in vivo

First, we tested whether IGF1-24 stimulated full-differentiated cardiomyocytes proliferation in vivo. We used 6-7-weeks old SD rats. At this age, cardiomyocytes have lost their proliferation ability[7]. Female SD rats were randomly divided into two groups, receiving either Phosphate-buffered saline (PBS) (control) or IGF1-24 daily for 7 days by tail intravenous injection at concentration of 100μg/kg/day (Fig2a). Cell proliferation was assessed by 5-ethynyl-2’-deoxyuridine (EdU) as well as proliferation antigen Ki-67 and histone H3 phosphorylated on serine10 (H3PS10). EdU is a uridine analogue that is incorporated into newly synthesized DNA[21]. For the EdU assessment, the rats were administered EdU intraperitoneally every 24h for additional 7 days (Fig2a). The percentage of EdU+ cardiomyocytes significantly increased up to 0.6% upon treatment with IGF1-24 injection (Fig2b). In agreement with this, treatment with IGF1-24 increased the number of the proliferation marker Ki-67(Fig2c) and H3PS10 compared to the control group (Fig2d). In addition, no changes were detected in IGF 1-24 treated hearts compared to controls in histological examination (Fig2e). Collectively, these results show that IGF1-24 stimulates endogenous cardiomyocyte proliferation in the healthy, uninjured rats.

**Fig.2.**
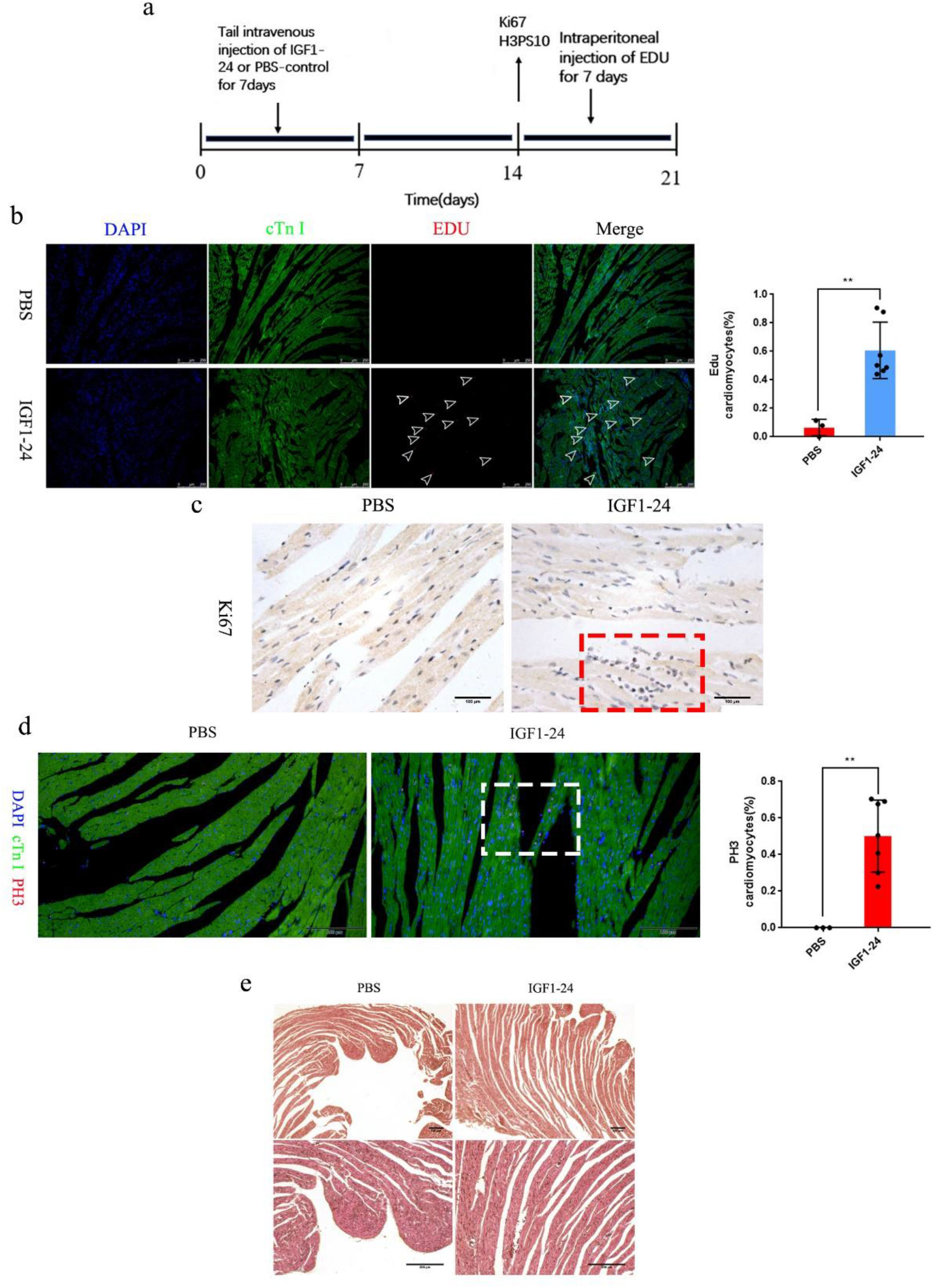
IGF1-24 treatment induces cardiomyocyte proliferation. **a,** Schematic representation of the protocol for IGF 1-24 administration in vivo. **b,** Representative images of EdU after IGF1-24 administration, with relative quantifications. The bottom images show higher magnification of the outlined regions in the top images. **c,** Representative images of cardiomyocytes with Ki-67+ nuclei immunostaining in the heart after IGF1-24 administration. **d,** Representative images of cardiomyocytes with H3PS10 nuclei immunostaining in the heart after IGF1-24 administration, with relative quantifications. **e**, HE staining of tissue sections after IGF1-24 administration. The bottom images show higher magnification of the outlined regions in the top images. Data are mean ± s.e.m.; the number of animals per group is indicated. Quantification is made from at three acquired from three different regions of each heart. *P < 0.05; Student’s two-sided t-test.

### IGF1-24 repaired cardiac injury in vivo after cardiac damage

Next, we tested the effects of IGF1-24 on cardiac repair in vivo after cardiac damage. Isoproterenol were used to induce myocardial injury[22, 23]. As a first step, 6-7-weeks old female SD rats were randomly divided into three groups. One group was administrated with PBS as control. Two groups were treated with intraperitoneal injection of isoproterenol(80mg/kg/day) daily for 7 days to induce myocardial injury, followed by administration of either IGF 1-24 (100μg/kg/day) (ISO+IGF1-24 group) or PBS (ISO group) by tail intravenous injection daily for another 7 days to the cardiac injured rats. For the assessment of EdU, rats in all groups were intraperitoneal injected EdU (total 50mg/kg) for additional 7 days (Fig3a). We observed obviously increased number of EdU+ cardiomyocytes in IGF1-24 treated group (1.52%) (Fig3b). In agreement with this, we also observed increases in the number of positive Ki-67 cardiomyocytes (Fig3c) and H3PS10 in IGF1-24 treatment group after cardiac injury (Fig3d). Furthermore, histological analysis showed an obvious improvement in heart tissue in IGF 1-24 treatment group compared to the isoproterenol treated group (Fig3e). Masson staining showed that IGF1-24 significantly reduced the degree of fibrosis after heart injury (Fig3f). Collectively, these data demonstrate that recombination protein IGF1-24 repaired myocardial injury by inducing cardiomyocyte proliferation.

**Fig.3.**
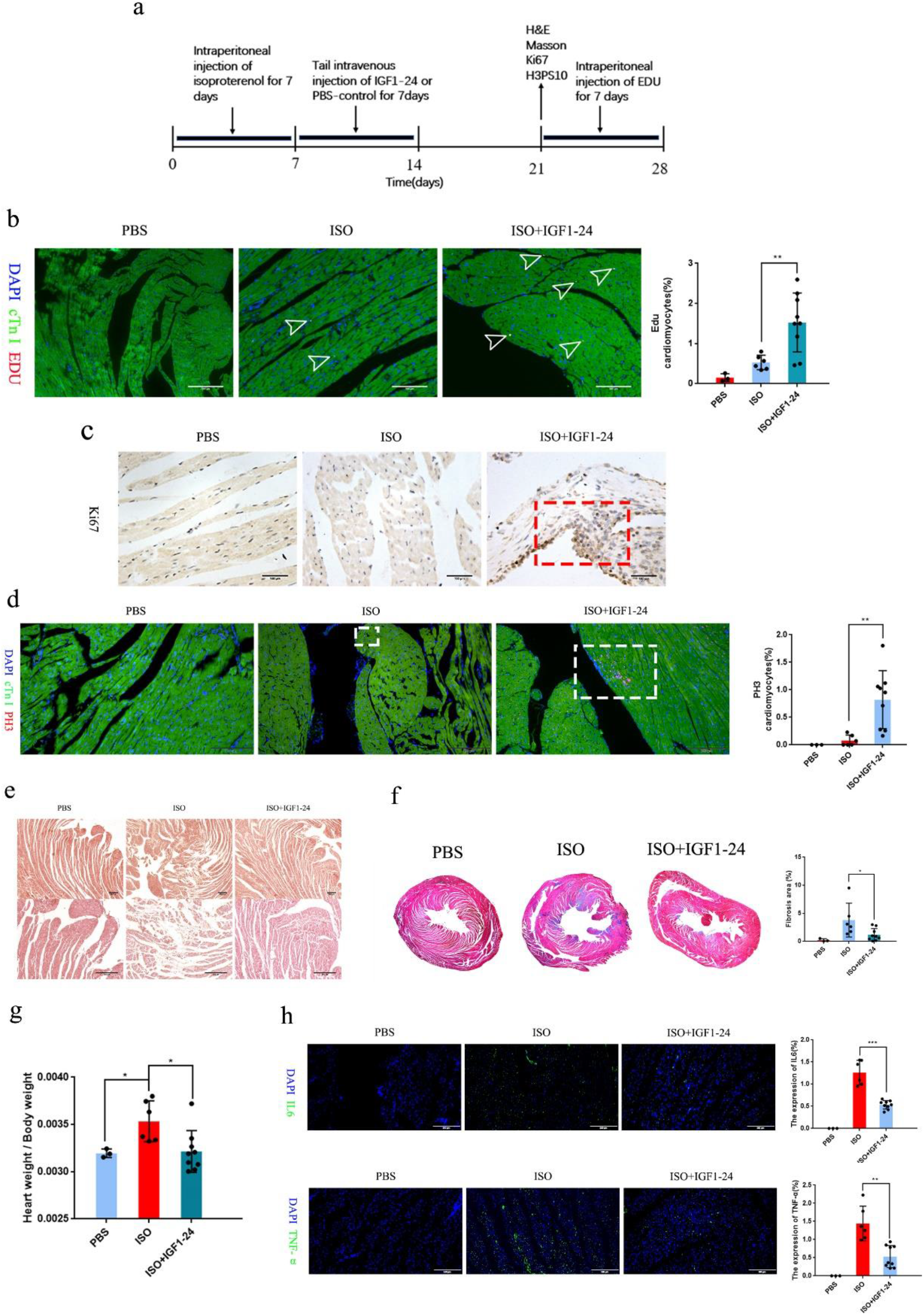
IGF1-24 administration improves cardicac repair. **a,** Damage model construction by subcutaneous injection with isoproterenol and Experimental protocol for EdU administration in vivo. **b,** Representative images of EdU in the injured heart after myocardial injury, with relative quantifications. **c,** Representative Ki-67+ immunofluorescence in the injured heart after myocardial injury. **d**, Representative H3PS10 immunofluorescence in the injured heart after myocardial injury, with relative quantifications **e,** HE staining of tissue sections from myocardial tissue after myocardial injury. The bottom images show higher magnification of the outlined regions in the top images. **f,** Masson trichrome staining of tissue sections from myocardial tissue after myocardial injury. **g**, statistics data of heart weight / body weight. **h**, Representative immunohistochemistry images of TNF-a, IL-6 in the injured heart after myocardial injury, with relative quantifications. Data are mean ± s.e.m.; the number of animals per group is indicated. Quantification is made from at three acquired from three different regions of each heart. *P < 0.05; Student’s two-sided t-test.

We also recorded the ratio of heart weights and body weights. The ratio increased significantly after injection of isoproterenol, however, treatment with IGF 1-24 returned the ratio toward the level of the control group (Fig3g). In addition, we observed significantly decrease of inflammatory factors (TNF-α & IL-6) (Fig3h) in IGF1-24 treatment group after cardiac infarction, indicating that lower inflammation happened after IGF 1-24 treatment.

## Discussion

Myocardial infarction can cause permanent loss of myocardial cells and the formation of scar tissue, leading to heart failure and irreparable damage to heart function. Cardiomyocytes are terminally differentiated cells and cannot proliferate and differentiate shortly after birth[24]. Therefore, it is urgent to find approaches and medicine that can effectively stimulate myocardial repair. For myocardial repair, there are two general directions so far: ① cell-based repair methods; ② factors related to the myocardial repair.

For the repair methods, non-cardiomyocytes[25, 26], cardiac-derived cells[27, 28], and pluripotent stem cells [29, 30]are differentiated into functional myocardial cells by transplantation in vivo, thereby improving the function of the heart. However, results showed that these cells rarely differentiate into cardiomyocytes in the body and may cause negative effects such as arrhythmia. Researchers also reported the reprogrammed non-cardiomyocytes in vitro [31, 32]and fibroblasts in vivo[33, 34], however, the reprogramming efficiency is low and it is not easy to deliver the reprogramming fibroblasts in vivo. In addition, regulating cell cycle has been reported to successfully stimulate cardiomyocytes re-enter the cell cycle and trigger proliferation[12], while endogenous stimulation exists the phenomenon of off-target cell proliferation.

For the factors related to the myocardial repair, several growth factors [35, 36] have been found to promote the proliferation of cardiomyocytes, but they have the risk to cause cardiac fibrosis. microRNAs [7,11] also show effects in stimulated cardiomyocytes proliferation in vitro and in vivo. When using microRNA therapy, large doses are required to maintain its concentration to prevent RNases from degrading it in the body[37]. The high level of microRNA may cause an immune response and thus cause an inflammatory response[38]. Exosomes[39, 40] are also used for myocardial repair, and there is a lack of effective local targeting to the site of action.

Here we designed the recombinant protein IGF 1-24 and investigated its effects on differentiated cardiomyctes in rats. We found that it triggered the myocardial cell proliferation at a higher than that reported[41]. Moreover, HE staining in rat heart tissues show that there is no difference in the cardiac tissue between the recombinant protein IGF1-24 group and the control group, implying that the recombinant protein IGF1-24 will not affect the healthy cardiac tissue. Next, we performed a model of myocardial injury by intraperitoneal injection of isoproterenol in rats. Then we Injected IGF1-24 to determine its effect on the damaged myocardial tissue repair. The results showed that the cardiomyocyte proliferation rate of the rats injected with IGF1-24 was 1.52%, which is much higher than other reports (the myocardial cell proliferation rates are 0.19% −0.9%)[7, 42, 43].

In conclusion, our research confirms that the recombinant protein IGF1-24 promotes the proliferation of cardiomyocytes, reduces the area of cardiac fibrosis, and reduces the levels of inflammatory factors. The IGF 1-24 could be a prospective medicine to heart repair because it has following advantages. (1) Instead of the comprehensive method of transplanting cells or transfection through vectors and in situ injection, the recombinant protein IGF1-24 can be intravenous injected into body. The application is very simple, providing a promising therapy for heart injury repair. (2) The effets of IGF 1-24 on cardiomyocye proliferation and heart regeneration are much higher than other reports.

## Materials and Methods

### Production and purification of recombinant IGF1-24

IGF 1-24 sequence was constructed by linking IGF1 and 24 amino acids which was cloned from C-terminal of IGF 1Ec by three glycine as shown in our previous report ^17^. IGF1-24 fragment was PCR-amplified introducing BamH I and Xho I sites, and then cloned in pET32a plasmid preserved in our laboratory, to generate a recombinant plasmid pET32a/IGF1-24, which was further confirmed by sequencing (Beijing Genomics Institute, China). Subsequently, pET32a/IGF1-24 was transformed into E.coil Rosetta (DE3) for recombinant protein IGF1-24 expression and purification. The protein expression in Rosetta(DE3) was induced by Isopropyl β -D-1-thiogalactopyranoside (IPTG 0.6mM) at 16°C for 6 h. Rosetta (DE3) pellets were collected by centrifugation and then ultrasonicated in ice-cold phosphate buffered saline (PBS) containing 1 mM DTT and protease inhibitors cocktail after triple freeze-thawed cycles. Cell lysate was centrifuged at 10000 rpm for 10 min at 4 °C and then the separated supernatant containing soluble IGF1-24 was incubated with Ni-Agarose and eluted by imidazole solution, that imidazole was finally removed by ultrafiltration. Recombinant protein IGF1-24 was dissolved in PBS.

### Injection of IGF1-24 in rats

Female Sprague-Dawley (SD) rats (6-7 weeks old), weighing 200g±20g, were purchased from Chongqing Medical University. The rats were divided to two groups by principle of randomness, one group was injected PBS (the same volume with IGF1-24 group) daily for 7 days as control and the other group was administrated with tail intravenous injection of IGF1-24(100μg/kg/day) daily for 7 days. At the end of studying, rats were anaesthetized and euthanized by abdominal injection of 10% pentobarbital sodium at concentration of 1ml /g. Weights of rats were weighed and recorded. Animal care and the protocol for the animal studies were performed in accordance with the regulations of the Animal Care and Use Committee at Army Medical University, China.

### Cardiac injury model and repair

Rat cardiac injury model was established by intraperiotoneal injections of isoproterenol^19,20^. Six-to-seven-week-old female SD rats, weighing 200g±20g, were injected isoproterenol at concentration of 80mg/kg/day every 24h for 7 days to induce cardiac injury. Then injured rats were randomly divided into two groups (ISO group and ISO+IGF1-24 group). Animals in ISO group were injected with PBS (the same volume with IGF1-24 group) daily for 7 days. Rats in ISO+IGF1-24 group were treated with IGF1-24 (100μg/kg/day) by tail intravenous injection every 24 hours for 7 days. Healthy rats without isoproterenol and/or IGF 1-24 treatment were injected with PBS and as control. At the end of study, rats were anaesthetized and euthanized by abdominal injection of 10% pentobarbital sodium at concentration of 1ml /g.

### Heart collection and histological analysis

Hearts were exposed, washed by PBS and weighed. Cardiac apexes were separated and dipped in 10% formalin at room temperature for overnight, then dehydrated by alcohol at increasing concentration gradient (50, 75, 95, 100%) for 60min. Cardiac apexes tissues were embedded in paraffin and cut into slices at 4 μm, dewaxed in xylene at twice for total 30 min and rehydrated in alcohols at decreasing concentration (100, 95, 75, 50%) at room temperature. Haematoxylin and eosin (H&E) was performed and analyzed following standard procedure.

### Immunofluorescence and immunohistochemistry analyses

In order to antigen retrieval, hydrated sections were boiled in 0.1M sodium citrate buffer at pH 6.0 by 20 min and cooled down at room temperature. All sections were washed by PBS for 3 min three times, permeabilized by 0.5% Triton X-100 for 30 min and blocked by 10% goat serum for 1 h at room temperature. Sections were stained with the following primary antibodies diluted in Bull Serum Albumin (BSA) overnight at 4°C, anti-cTnI (Proteintech), anti-Ki-67(BIOSS), anti-H3PS10(ZEN BIO),anti-TNF-α (ZEN BIO), anti-IL-6(ZEN BIO). Sections were washed by PBS for 3 min three times and incubated respectively with the following secondary antibodies, FITC-labeled Goat Anti-Rabbit IgG(H+L) (Beyotime), Cy3-labeled Goat Anti-Rabbit IgG(H+L) (Beyotime) and FITC-labeled Goat Ani-Mouse IgG(H+L) (Beyotime) for 60min.Nuclei were stained with DAPI (Leagene). Alternatively, after endogenous peroxidase was inhibited by 3% H2O2, sections were permeabilized by 0.5% Triton X-100 for 30 min and blocked by 10% goat serum for 1 h at room temperature. Sections were stained with antibodies overnight. After washed by PBS, secondary antibodies were stained on sections for 60 min, DAB solution was stained in sections for 1 min. Sections were observed in 500× magnification. Immunofluorescence images were analyzed by Image J.

### EdU assay

To detect cardiomyocyte proliferation in vivo, EdU (total 50mg/kg) was injected intraperitoneally to rats for 7 days. At the end of the study, rats were anaesthetized and euthanized. The excised hearts were washed by PBS, weighted and frozen at liquid nitrogen. For EdU analysis, sections were permeabilized. According to operation manual, reaction buffer was mixed. Sections were stained with reaction buffer for 30 min. All sections were washed by PBS for 3 min three times. Sections were observed in 100× and 200× magnification, and at least 3 sectors for each sections. Images were analyzed by Image J.

### Masson staining

The sample sections were dewaxed and hydrated by conventional methods, stained with weigert hematoxylin for 5min, differentiated by acidic ethanol for 10s, and thoroughly cleaned by water. Sections were stained with ponceau for 10min and cleaned with acetic acid solution. Then sections were treated phosphomolybdic acid solution for 30s, was re-dyed with aniline blue solution for 30s and treated with acetic acid solution until no blue peel was found. The sample sections were dehydrated, transparent and sealed with gum, then observed and photographed under an optical microscope. The degree of fibrosis was calculated with Image J.

### Statistical analysis

Data are presented as mean ± s.e.m and analyzed by Student’s two-sided t-test. The statistical graphs are made by graph pad. All statistical analyses, significance was controlled at P < 0.05.

## Author contributions

L.T. designed the experiments and supervised the project. X. W., M.F., Y. L., and Y. C., performed the in vivo experiments. Q.Y and J.F. designed the sequence and purified the recombinant protein IGF 1-24. X.X. optimized the separation and purification of IGF 1-24. X.W., M.F., and P.L. performed the histological and immunohistochemistry analyses. M.F., X. W.,and Y. L. performed the data analysis and statistic. P.L., H.S. and Y.L, performed the evaluation of rats conditions. M.F., X. W., Q.Y., J.F. and Y. L. wrote the manuscript.

## Supplementary Materials

**Figure S1.**
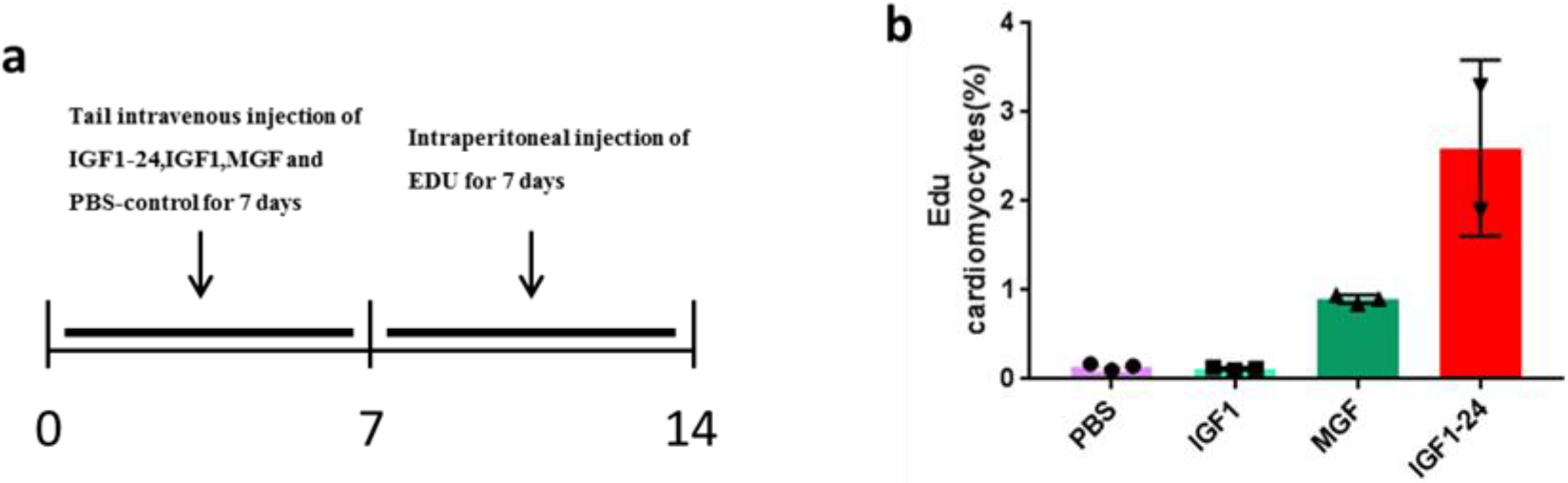
IGF1-24 treatment induces cardiomyocyte proliferation. **a**,Schematic representation of the protocol for IGF 1-24 administration in vivo. **b**, Statistics of Edu proliferation results in different groups.

